# From receptor binding to biogeography: Multi-scale prediction of filovirus hosts in bats

**DOI:** 10.64898/2026.05.18.726005

**Authors:** Adrian A. Castellanos, Simon J. Anthony, Kartik Chandran, Gorka Lasso, Heather L. Wells, Barbara A. Han

## Abstract

Forecasting zoonotic risk requires identifying which host species are biologically susceptible to infection, yet susceptibility is rarely predicted using frameworks that integrate molecular mechanisms with macroecology. Filoviruses, a diverse group of bat-associated viruses that include Ebola and Marburg viruses, illustrate this challenge: viral entry depends on interactions between viral glycoproteins and the host receptor NPC1, and host ecology and distribution determine opportunity of viral entry. Additionally, receptor sequence data used for informing viral entry are available for only a small fraction of bat species. Here, we extend virus-specific susceptibility prediction across the global diversity of bats by integrating experimentally measured and physicochemically inferred virus–receptor binding strengths with phylogenetic, ecological, and environmental data. Using boosted regression models trained on binding assay labels, we generate predictions of NPC1-mediated binding strength for more than 1,300 bat species. Predicted susceptibility is strongly structured by evolutionary relationships, with high binding concentrated in particular bat lineages, but is further differentiated within clades by morphology, life-history strategy, and environmental context. Strikingly, macroevolutionary structure alone recovers interaction patterns originally derived from amino acid–level physicochemistry, indicating that information about receptor-mediated compatibility is recoverable from host evolutionary history and ecological traits. Predicted high binding strength extends well beyond historically recognized outbreak regions, suggesting that the fundamental host range of filoviruses may be substantially broader than their currently realized distribution. By scaling receptor biology to global host diversity, this multi-scale framework expands mechanistic susceptibility forecasting beyond species with available molecular data and provides a generalizable approach for integrating molecular and ecological information in zoonotic prediction.

## INTRODUCTION

Predicting spillover transmission is a central goal in disease ecology. When made with sufficient precision and mechanistic understanding, predictions about animal hosts, locations, and timing of spillover events can enable upstream interventions that reduce zoonotic disease risk at its source. Achieving this level of predictive resolution, however, remains challenging for pathogens with broad host ranges and complex ecological contexts.

Recent years have seen substantial advances in upstream prediction of zoonotic hosts, driven by trait-based and phylogenetically informed models applied across broad taxonomic groups and pathogen systems. These approaches have proven useful for identifying likely hosts and prioritizing surveillance, both for zoonoses broadly [1–3] and for targeted pathogen groups. For instance, host predictions have been made for filoviruses [4,5], henipaviruses [6], coronaviruses [7], leishmaniasis [8], *Borrelia burgdorferi [9]*, Zika virus [10], rabies virus [11], orthopoxviruses [12], and mammarenaviruses and hepaciviruses [13].

Despite these advances, many of these predictions still remain too coarse to directly inform intervention. Trait-based models often identify large sets of plausible hosts without resolving which species are likely to maintain particular pathogen species, or under what ecological conditions spillover risk may increase. This lack of specificity becomes especially problematic for pathogen groups with many closely related viruses and large, ecologically diverse host clades.

Filoviruses illustrate this challenge well. Of the eleven recognized mammalian filoviruses, only five are known to have caused human outbreaks (Taï Forest causes human disease but only has one documented case, [14] Figure 1), yet all are suspected to circulate in sylvatic reservoirs between outbreaks [15]. Bats are widely implicated as reservoir hosts [15,16], but this inference spans a clade of roughly 1,500 species with extraordinary ecological, physiological, and biogeographic diversity ([17,18]).

**Figure 1.**
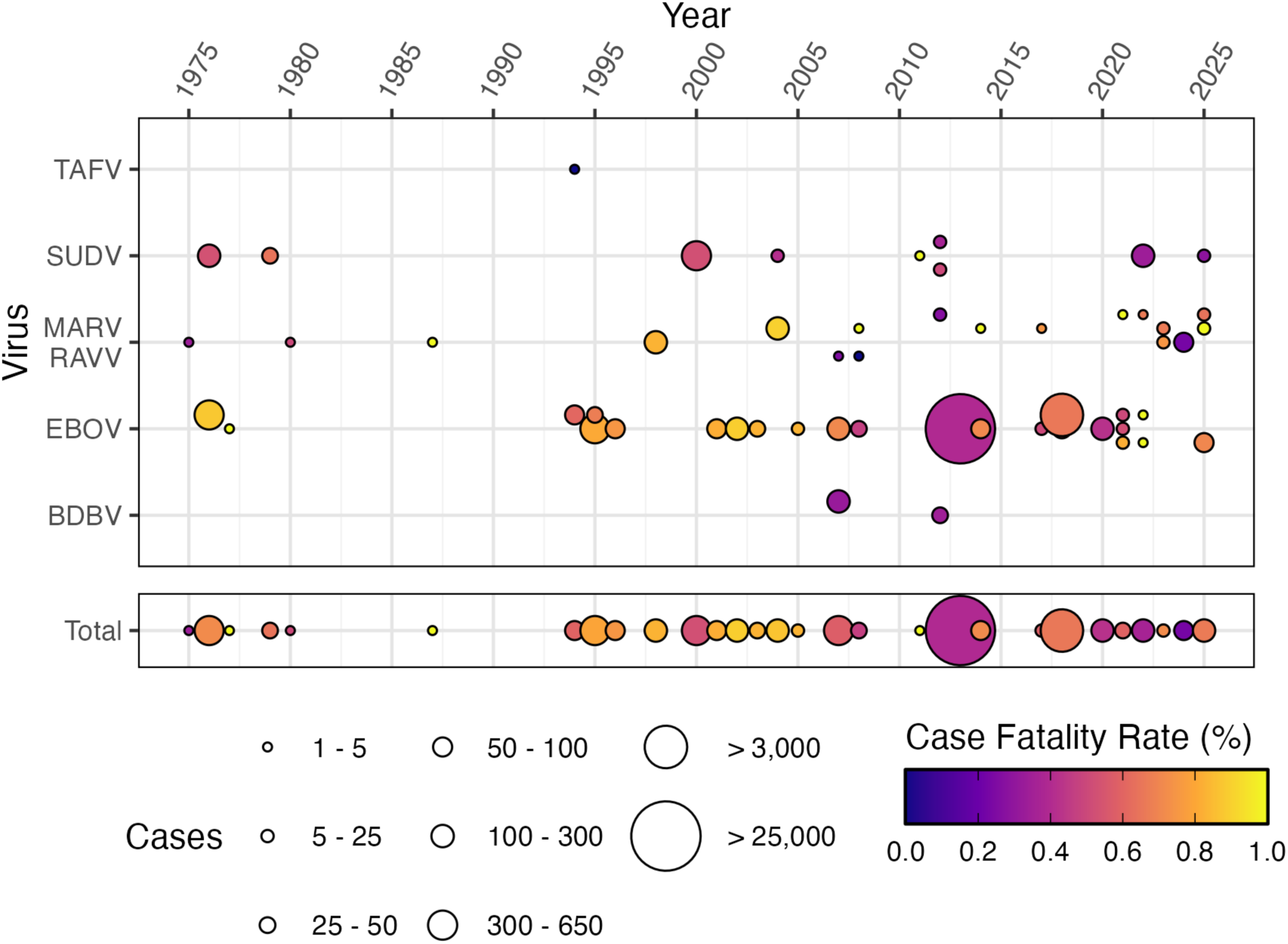
Plot showing documented filovirus outbreaks. The x-axis documents time in years, while the y-axis depicts the filovirus responsible for the outbreak. MARV and RAVV are combined as they both cause Marburg virus disease. The size of the circle represents the size of the outbreak (number of cases), while its color relates the reported case fatality rate (deaths/cases) with lighter colors showing a higher case fatality rate. Note that some years have multiple documented outbreaks. The bottom section represents the total disease burden caused by filoviruses in each year.

**Figure 2.**
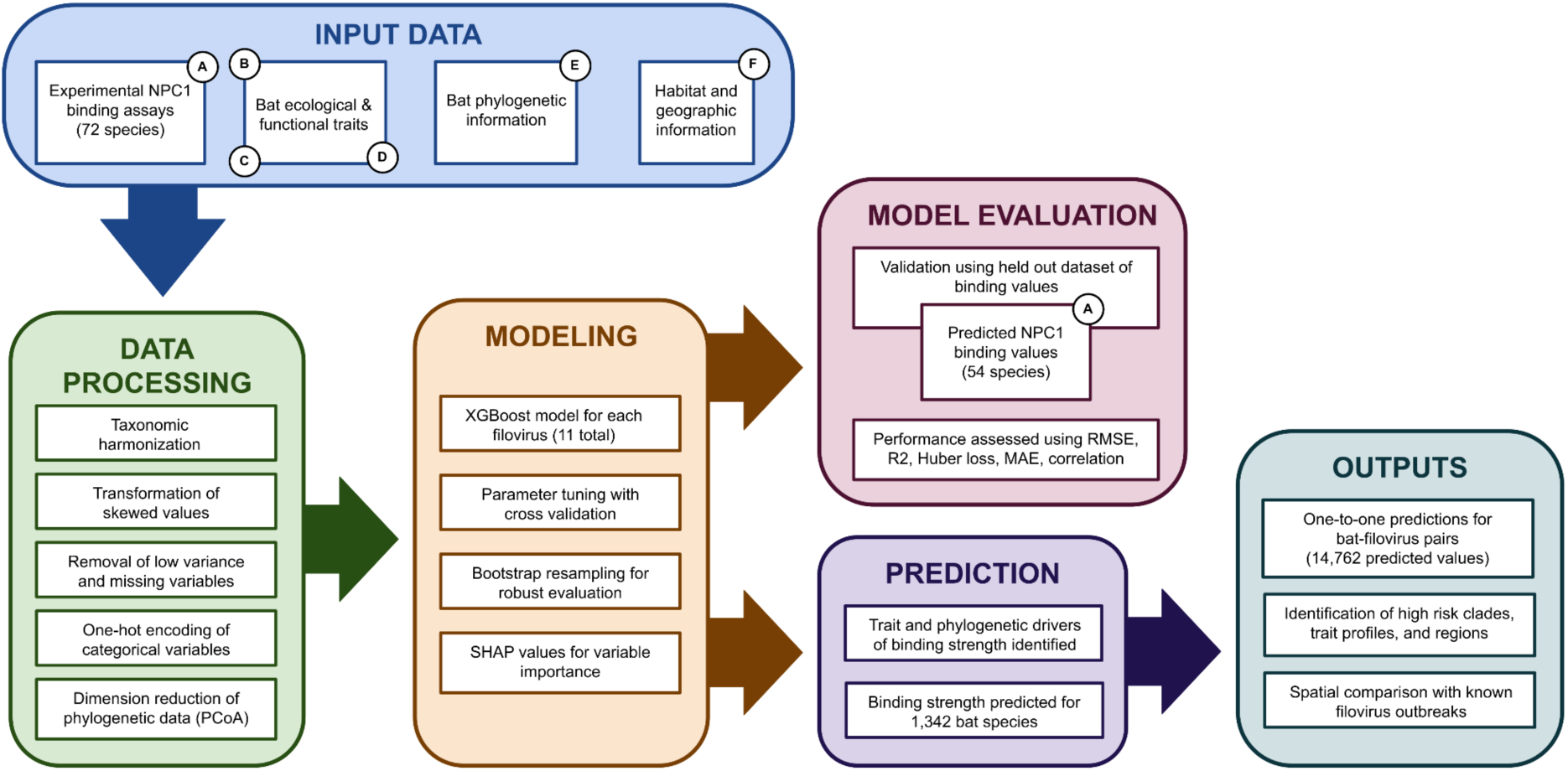
Each colored box represents a step in our pipeline that contains multiple elements. Arrows show the flow of data through the pipeline. References to training and label data are given as circles: A) Lasso et al. 2025 B) Soria et al. 2021, C) Crane et al. 2022, D) Guy et al. 2020, E) Upham et al. 2019, F) IUCN Red List 2023.

Among the bat species overlapping filovirus-endemic regions, evidence for susceptibility varies widely and is often context dependent. Laboratory and field studies report heterogeneous patterns of seropositivity and experimental infection, including apparent resistance in some species (e.g., *Eidolon helvum* [19]), alongside positive detections in others (e.g., *Mops condylurus*, *Miniopterus schreibersii*, *Rousettus aegyptiacus*, *Rousettus leschenaultii* [20–23]). In natural systems, infection susceptibility is further shaped by environmental stressors that are difficult to measure at scale, including seasonal resource limitation ([24,25]), reproduction ([26,27]), and patterns of land-use change that are historically dynamic and ongoing ([28,29]). For widely distributed species, the spatial and temporal variability in these stressors increases uncertainty about when and where surveillance is most appropriate [30].

Incomplete sampling and uneven ecological and taxonomic data further complicate the identification of likely host species. Patterns of cryptic diversity within climate refugia that overlap filovirus hotspots vary across bat lineages and traits [31,32], and many undersampled species are consequently excluded from predictive analyses due to insufficient ecological and genetic data. At the same time, bat movements and population connectivity vary substantially, from cave-dependent species with fragmented or weakly connected subpopulations to long-distance fliers capable of traversing hundreds of kilometers in a single foraging bout [33]. Together, these factors make surveillance logistically challenging and difficult to justify when predictions cannot distinguish between bats likely to harbor zoonotic filoviruses (e.g., Ebola virus) from those associated with filoviruses not currently recognized as spillover threats.

Initial efforts to predict filovirus hosts collated species-level trait data for a subset of bat species and combined these with generic filovirus seropositivity records, which are typically not resolved to individual filovirus species [34], to generate coarse predictions of potential reservoir hosts [4]. Despite the coarseness of this approach, several high-ranking predictions were later corroborated by field studies, including detections of filovirus exposure in previously unsampled species [22,35] and the discovery of novel filoviruses in hosts whose trait profiles placed them among the most likely reservoirs (Bombali virus; [20]). These successes underscored both the utility and the limitations of trait-based prediction: such approaches can identify host species that warrant surveillance, but they are less reliable for resolving which virus species are most likely to occur in which hosts. This lack of *dual precision* in host-virus pairs is particularly important when host range or viral diversity are high, characteristics that are shared among the most consequential contemporary zoonotic threats (e.g., SARS-CoV-2, filoviruses, avian influenza).

Subsequent work addressed these limitations in two ways: by using more consistent molecular proxies of susceptibility, and by increasing predictive resolution from pathogen families to individual virus species. Working with the SARS-CoV-2 Wuhan-1 variant, computational estimates of molecular binding strength reduced reliance on geographically and taxonomically biased surveillance data and enabled predictions about an emerging pathogen [36]. For filoviruses, molecular binding assays quantified one-to-one interactions between the filovirus-specific glycoproteins and Niemann-Pick C1 (NPC1) receptor orthologs of multiple host species, revealing physicochemical determinants of viral entry across host-virus pairs [37]. These approaches developed a mechanistic foundation for predicting host susceptibility to specific viruses that complements trait-based ecological inference by anchoring broad ecological patterns in the molecular interactions that are necessary for infection.

While powerful, this strategy remains fundamentally constrained by data availability. Reliable NPC1 ortholog sequences exist for only a fraction of bat species, restricting empirical screening to a narrow subset of a large and diverse clade. Of approximately 1,500 bat species, NPC1 sequences in Lasso et al. [37] were available for only 122 unique species. In practice, this means that the host-virus associations that can be characterized most precisely are often those drawn from a limited, unevenly sampled portion of the bat phylogeny, leaving large geographic regions and taxonomic groups effectively unexamined. These limitations notwithstanding, Lasso et al. [37] represents the most comprehensive empirical analysis of filovirus receptor compatibility to date, but predictions remain bounded by uneven and limited availability of basic molecular data, and limited integration of ecological context.

An open question is whether spillover risk will remain low for some filoviruses—perhaps due to intrinsic incompatibility with human hosts—or instead increase for viruses that have historically been quiescent under changing environmental conditions. Filoviruses such as Lloviu virus (LLOV) and Taï Forest virus (TAFV), have not caused large outbreaks in human populations, but show varying evidence of their ability to infect humans with TAFV being responsible for one human case [38] and LLOV showing the ability to infect non-human primate cells [39,40]. With LLOV now identified in Spain, Hungary, Italy, Bosnia and Herzegovina, and China [21,39,41–43], and with evidence for human infectivity across these understudied filoviruses remaining uneven, these patterns illustrate the uncertainty surrounding zoonotic potential in systems with broad host ranges and wide geographic distributions. At the same time, recent human outbreaks of Marburg virus (MARV) and Sudan virus (SUDV; Figure 1), both previously considered rare causes of spillover infection, underscore how rapidly risk profiles can shift. Anticipating spillover in this dynamic landscape requires predictions that are both mechanistically grounded and sufficiently resolved across host diversity to be operationally useful.

Here, we present a framework for achieving dual precision in filovirus prediction: resolving which bat species are likely hosts for which filovirus species. Across all eleven bat-associated filoviruses, we generate one-to-one predictions over the full diversity of bat species by integrating experimental and inferred molecular binding information with host traits that capture ecological and evolutionary context. By augmenting sparse NPC1 receptor sequence data using trait-based inference, this multi-scale fusion approach extends mechanistic prediction beyond the limits imposed by uneven molecular sampling. In doing so, it advances predictive capacity for Ebola and related filoviruses while also exposing fundamental knowledge gaps where data augmentation cannot substitute for missing biological understanding.

Our results corroborate scant experimental evidence to suggest that susceptibility to filoviruses, as inferred from NPC1 binding, is widespread across the bat phylogeny, encompassing unexpected families and species occurring well outside the currently recognized geographic distribution of filoviruses. The broad spatial pattern of predicted susceptibility likely reflects underlying phylogenetic structure, with additional contributions from species-level trait profiles shaped by geography, morphology, and life history. Beyond refining host prediction, this approach generates new hypotheses about filovirus biogeography and co-evolutionary history with bat hosts, and highlights data gaps where additional empirical work would most improve predictive resolution. Although demonstrated here for filoviruses, this multi-scale integration of molecular, evolutionary, and ecological information provides a generalizable template for improving predictions for zoonotic pathogens with similarly broad host ranges, offering a scalable path toward the dual precision needed to anticipate and prevent spillover at its source.

## RESULTS

### Trait data

We collected data on 119 trait variables, informing phylogenetic position, diet, habitat preference, reproductive characteristics, morphology, geographic location, and dispersal ability. Of these, we retained 76 variables after removing those showing near zero variation (37 variables removed, mainly uninformative habitat or locomotion characteristics) or those missing more than 30% of their values in the training data (six additional variables removed).

### Boosted regression

Boosted regression models predicting NPC1 binding strength across 11 filoviruses showed variable performance depending on the evaluation metric and dataset (cross validation test data vs. held out validation data), with the majority of models demonstrating reasonable to high levels of support (Tables 1 and 2). With the exception of MLAV, the *Orthomarburgvirus*-like filoviruses (DEHV, MARV, RAVV) show among the best evaluation metrics in the validation dataset with Huber loss values under 30 and R^2^ values above 0.44. Of the *Orthoebolavirus*-like filoviruses, BDBV, BOMV, EBOV, and LLOV showed the best performance metrics, with validation Huber loss values under 40 and R^2^ values above 0.32. Intriguingly, SUDV and TAFV both display low Huber loss and RMSE values but also low R^2^ and correlation coefficients (Table 2). Of these, SUDV shows poor evaluation metrics when evaluated using cross validation. Both SUDV and TAFV also show the highest mean observed binding strength and most skewed binding strength distributions (−0.948 for TAFV and −1.488 for SUDV). In contrast, RESTV shows high cross-validation R^2^ values and strong correlation with observed values in the validation dataset, but poor validation performance and among the highest RMSE values across evaluation datasets.

**Table 1.** Results from the parameterization and evaluation of the XGBoost models created for each filovirus. Columns include values for the 7 tuned parameters (eta, max depth, subsample ratio, columns sampled by tree, minimum child weight, gamma, number of trees) and the evaluation metrics (RMSE and R^2^) for each model on training (using four-fold cross validation) and held-out testing data. EBOV - Ebola virus, BOMV - Bombali virus, BDBV - Bundibugyo virus, DEHV - Dehong virus, TAFV - Tai Forest virus, RESTV - Reston virus, SUDV - Sudan virus, MLAV - Měnglà virus, LLOV - Lloviu virus, MARV - Marburg virus, RAVV - Ravn virus. Binding scores ranged from 11 to 405.

| Filovirus | eta | Max depth | Subsample ratio | Columns sampled by tree | Min. child weight | gamma | Number of trees | RMSE |  |  |  | R<sup>2</sup> |  |
| --- | --- | --- | --- | --- | --- | --- | --- | --- | --- | --- | --- | --- | --- |
|  |  |  |  |  |  |  |  | Train | Train(%) | Test | Test (%) | Train | Test |
| **BDBV** | 0.00561 | 12 | 0.522 | 0.355 | 10 | 3.54 | 1,847 | 66.0 | 18.0 | 64.0 | 17.4 | 0.239 | 0.246 |
| **BOMV** | 0.00561 | 12 | 0.522 | 0.355 | 10 | 3.54 | 1,847 | 49.1 | 20.0 | 53.0 | 21.6 | 0.127 | 0.136 |
| **DEHV** | 0.0549 | 11 | 0.945 | 0.529 | 10 | 6.42e-6 | 9,801 | 35.7 | 14.6 | 34.0 | 13.8 | 0.510 | 0.532 |
| **EBOV** | 0.00561 | 12 | 0.522 | 0.355 | 10 | 3.54 | 1,847 | 60.4 | 17.6 | 57.6 | 16.7 | 0.423 | 0.445 |
| **LLOV** | 0.186 | 11 | 0.954 | 0.914 | 12 | 0.0124 | 7,848 | 43.6 | 15.4 | 41.6 | 14.7 | 0.486 | 0.516 |
| **MARV** | 0.0549 | 11 | 0.945 | 0.529 | 10 | 6.42e-6 | 9,801 | 35.3 | 14.4 | 33.1 | 13.5 | 0.468 | 0.494 |
| **MLAV** | 0.117 | 6 | 0.572 | 0.842 | 17 | 7.33e-9 | 3,692 | 74.4 | 22.2 | 84.3 | 25.1 | 0.189 | 0.127 |
| **RAVV** | 0.0549 | 11 | 0.945 | 0.529 | 10 | 6.42e-6 | 9,801 | 35.3 | 14.4 | 33.1 | 13.5 | 0.470 | 0.496 |
| **RESTV** | 0.00289 | 12 | 0.918 | 0.529 | 10 | 0.231 | 8,094 | 74.9 | 19.8 | 69.1 | 18.3 | 0.418 | 0.464 |
| **SUDV** | 0.00561 | 12 | 0.522 | 0.355 | 10 | 3.54 | 1,847 | 59.9 | 16.2 | 80.0 | 21.6 | 0.110 | 0.050 |
| **TAFV** | 0.00561 | 12 | 0.522 | 0.522 | 10 | 3.54 | 1,847 | 46.1 | 14.0 | 45.3 | 13.8 | 0.337 | 0.313 |

**Table 2.** Results from the comparison of predicted and observed values using held out validation data. EBOV - Ebola virus, BOMV - Bombali virus, BDBV - Bundibugyo virus, DEHV - Dehong virus, TAFV - Tai Forest virus, RESTV - Reston virus, SUDV - Sudan virus, MLAV - Měnglà virus, LLOV - Lloviu virus, MARV - Marburg virus, RAVV - Ravn virus.

| **Model** | **RMSE** | **R<sup>2</sup>** | **MAE** | **Huber**<br>**loss** | **Spearman** | **Pearson** |
| --- | --- | --- | --- | --- | --- | --- |
| BOMV | 30.548 | 0.344 | 23.687 | 23.211 | 0.485 | 0.587 |
| BDBV | 51.089 | 0.372 | 39.340 | 38.843 | 0.469 | 0.610 |
| EBOV | 54.446 | 0.326 | 40.198 | 39.698 | 0.496 | 0.571 |
| LLOV | 48.157 | 0.344 | 35.035 | 34.537 | 0.585 | 0.586 |
| RESTV | 65.157 | 0.296 | 49.311 | 48.811 | 0.486 | 0.544 |
| SUDV | 39.341 | 0.000 | 24.420 | 23.929 | 0.181 | -0.010 |
| TAFV | 36.800 | 0.129 | 29.423 | 33.313 | 0.387 | 0.360 |
| DEHV | 36.843 | 0.534 | 28.254 | 27.754 | 0.667 | 0.731 |
| MARV | 37.139 | 0.443 | 28.514 | 28.015 | 0.615 | 0.666 |
| MLAV | 69.863 | 0.292 | 57.285 | 56.785 | 0.548 | 0.541 |
| RAVV | 36.635 | 0.456 | 28.343 | 27.844 | 0.625 | 0.675 |

Looking at the individual predictions explains some of the patterns observed in our summary evaluation metrics (S2 Figure). Outliers, defined as species with an absolute difference between predicted and observed binding strength greater than 50, range from six (BOMV, SUDV) to 24 (RESTV) with a median number of 11 outliers per model. These outlier species are not consistently identified across models, with the exception of *Miniopterus minor*, *M*. *natalensis*, *Hipposideros lekaguli*, *H. diadema*, *Mops nanulus*, and *Rhinolophus ferrumequinum*. Some of these outlier patterns likely reflect limited sample size in our training dataset, particularly the absence of phylogenetic information from representatives from Noctilionidae and Miniopteridae. These individual predictions also reveal that *Orthomarburgvirus*-like filovirus models show stronger concordance between predicted and observed values for individuals in our training dataset (mean difference = 0.804) compared to *Orthoebolavirus*-like filoviruses (mean difference = 14.169).

SHAP values revealed consistent patterns in the importance of ecological and phylogenetic predictors of filovirus binding strength. Across models, a large proportion of the most influential variables encoded phylogenetic information from PCoA axes, comprising 36.8% to 57.9% of the top 19 variables, with a median of 47.4%. This pattern is driven in part by the shared importance of seven of the 17 PCo axes, which showed mean SHAP values greater than three for at least 10 models (Figure 3A). These same variables also showed similar patterns of marginal effect across models, indicating that similar taxa were consistently associated with higher predicted binding strength, albeit with varying magnitude (Figure 3B). Collectively, SHAP values identified bats in the families Molossidae, Pteropodidae, and Hipposideridae as having higher than expected binding strength, particularly along PCo axes 4, 8, 15, and 17.

**Figure 3.**
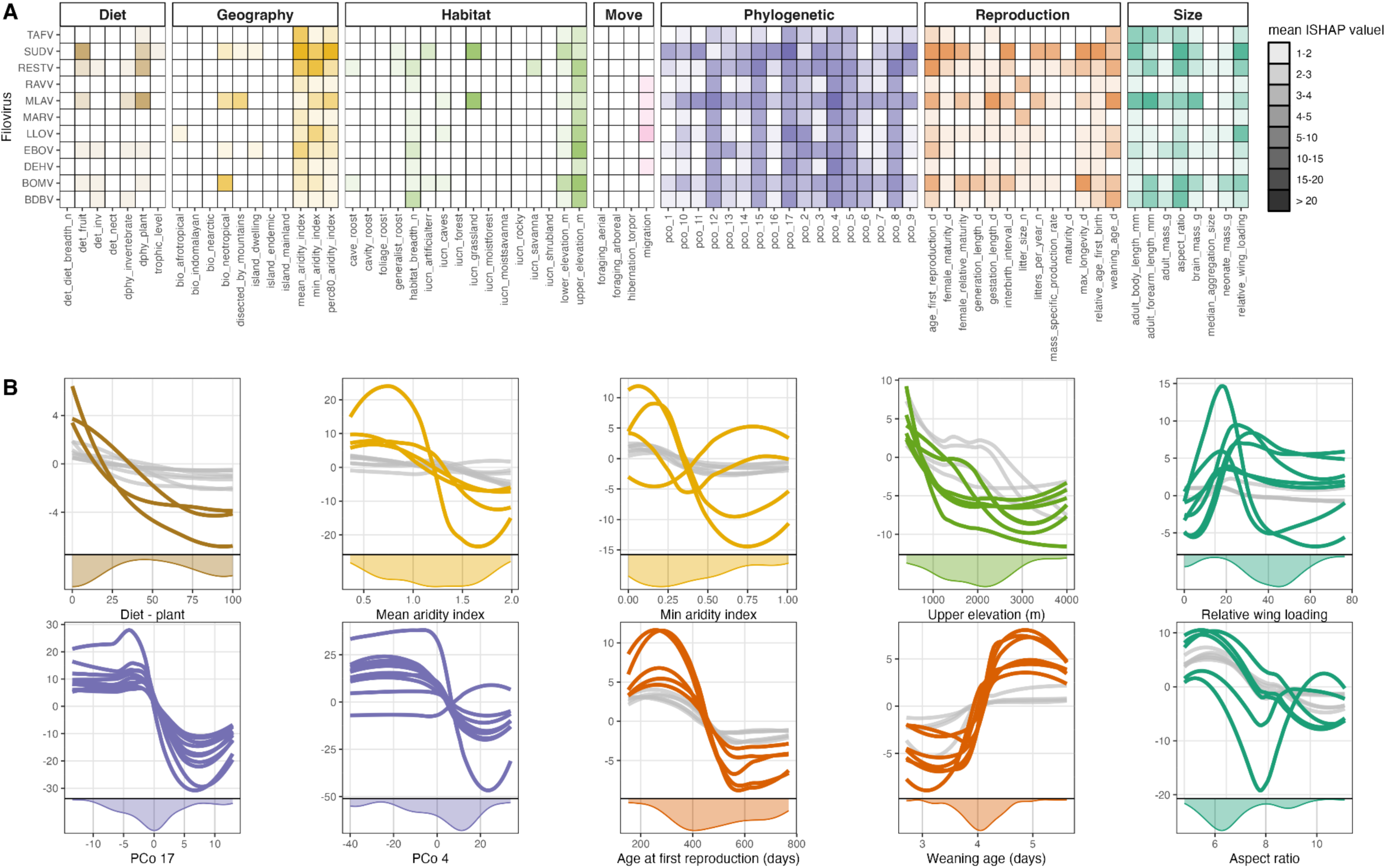
A comparison of variable importance on prediction and their marginal effects on prediction accuracy. The top panel shows the variable importance by mean absolute SHAP value for each variable and filovirus. Variables are organized and colored by type and the darker the color, the greater the importance. The 10 panels below show partial dependence plots for 10 variables for 11 filoviruses. Colored lines denote mean absolute SHAP value > 3, gray lines are SHAP values < 3.

The quantitative nature of these variables enabled further resolution at the genus and subfamily level rather than only at the family level. For example, PCo4 highlights most pteropodids, molossids, and select subfamilies of vespertilionids (S3 Figure). In contrast, PCo17 shows higher binding strength for select pteropodids (e.g., Epomophorinae) while showing substantial variation among hipposiderids (e.g., *Triaenops* vs. *Asellia*) and *Rhinolophus* (S4 Figure). The most speciose molossid genera and *Rousettus* are consistently associated with multiple influential axes, whereas other taxa with higher predicted binding strength were only linked to one (*Acerodon*, *Cynopterus, Pteropus*) or two (*Dobsonia*, *Nyctimene*, *Pteralopex*) axes (S3-6 Figures).

Although phylogenetic variables drive a substantial portion of our predictions, important variation is captured by functional traits and geographic variables as identified by SHAP values. Among these, the most important predictors typically describe aridity, reproductive characteristics, and morphology, whereas variables describing diet, habitat associations, movement, and broader geographical descriptors are generally uninformative (Figure 3A).

Based on absolute SHAP values, which summarize the magnitude of a variable’s effect on prediction regardless of direction, 34 non-phylogenetic variables (57.6%) show mean absolute SHAP values below five across models, while no phylogenetic variable falls below this threshold. The remaining informative non-phylogenetic predictors exhibit greater variability in both directionality and magnitude across models compared to the more consistent effects associated with the phylogenetic PCo axes (Figure 3B).

A notable exception to these broad patterns was the signal associated with elevation and aridity. Across most filoviruses, predicted binding strength was lower in bat species associated with higher elevations and higher in species associated with more arid conditions. This aridity effect was less consistent, however, for DEHV, MARV, RAVV, and RESTV when aridity was summarized using the 80th percentile of the aridity index, defined here as the value below which 80% of a species observed aridity values fall. Thus, low values of this metric indicate that most observations came from highly arid places and times. For these viruses, species with low aridity values at the 80th percentile instead showed lower predicted binding strength, contrary to the broader pattern. One possible explanation is that this metric becomes decoupled from other summaries of aridity, such as the minimum or the mean, in wide ranging species that occupy more heterogeneous climatic conditions. However, support for that explanation was mixed: the aridity index was only weakly correlated with range size (r = 0.15) and negatively correlated with habitat breadth.

We also saw convergence across models for variables describing bat size and reproductive characteristics. Higher binding strength was generally associated with earlier age at first reproduction and lower relative wing loading (RWL), which is a morphological estimation of wing area compared to body weight that is scaled by body mass in which a lower RWL represents a larger wing area independent of body mass. Several filoviruses (BDBV, EBOV, RESTV, and TAFV) were associated with larger species (longer length of body and forearm, larger aspect ratio, higher brain mass). DEHV, MARV, and RAVV showed higher binding strength for species with lower litter sizes. BDBV, EBOV, RESTV, and TAFV were associated with longer weaning ages. Although diet and dispersal traits are not well represented among important variables, higher binding strength was associated with lower plant contribution to the diet in RESTV and for migratory bats in LLOV, MARV, and RAVV.

We predicted binding strength for 1,342 bat species across 11 filoviruses, yielding 14,762 predictions (S7 Table). Predicted binding strength values ranged from 3.226 (*Rhinolophus ferrumequinum* to MLAV) to 414.817 (*Pteronotus rubiginosus* to MLAV). Mean predicted values ranged from 144.128 (BOMV) to 322.697 (SUDV), with SUDV and TAFV showing higher predicted binding strength in concordance with the observed binding strength values used to train these models.

Consistent with the strong influence of phylogenetic predictors, the highest binding strength predictions clustered in particular clades (Figure 4, S8 Figure). In particular, Molossidae, Pteropodidae, and Hipposideridae showed the highest predicted binding strength across models, regardless of the filovirus (Figure 4). Several vespertilionid species also exhibited high binding strength (S8 Figure), which is notable given their high species richness in the Americas and limited prior association with filoviruses.

**Figure 4.**
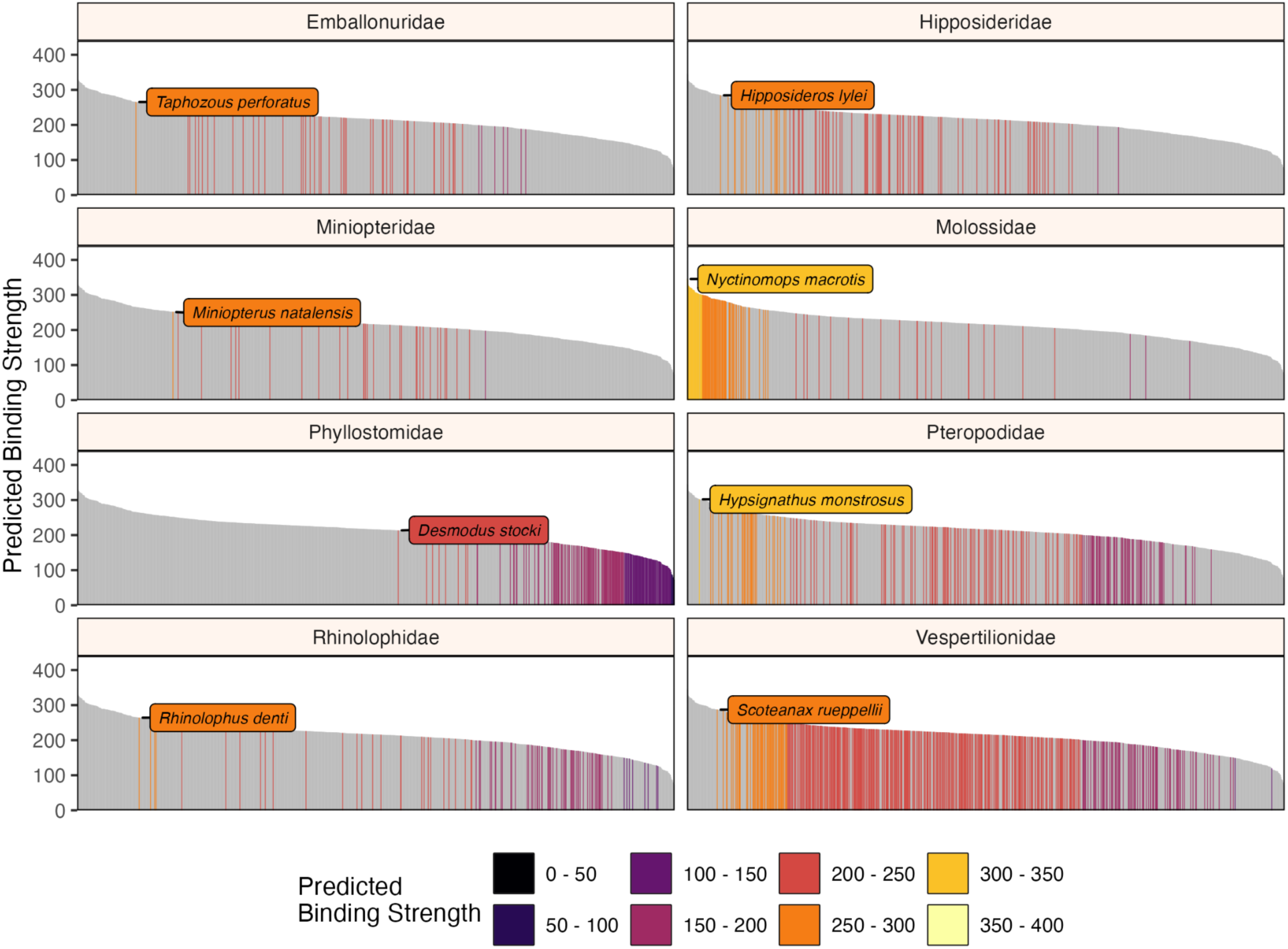
Model predictions presented for ebolavirus (EBOV) across all bats shown for the eight most speciose bat families. The y-axis shows predicted binding strength to EBOV for each bat NPC1, while the x-axis arranges the bat species in order from highest binding to lowest binding. Each panel depicting a bat family shows only those species in the family with the color of the bar relating to a binned binding strength category. The species with the highest predicted binding strength for each family is labeled.

Predicted high-binding species spanned all geographic regions, despite the inclusion of broad variables describing biogeographic realms that were largely unimportant for all models. Their distributions were not restricted to regions with documented outbreaks or detections, including sub Saharan Africa, parts of China, and Europe. Geometric mean of binding strength mapped onto bat distributions showed that Africa and southeastern Asia have the highest binding strength for most viruses, but additional pockets of high predicted binding strength were identified in South America, Mexico, Australia, and several regions of Europe (Figure 5).

**Figure 5.**
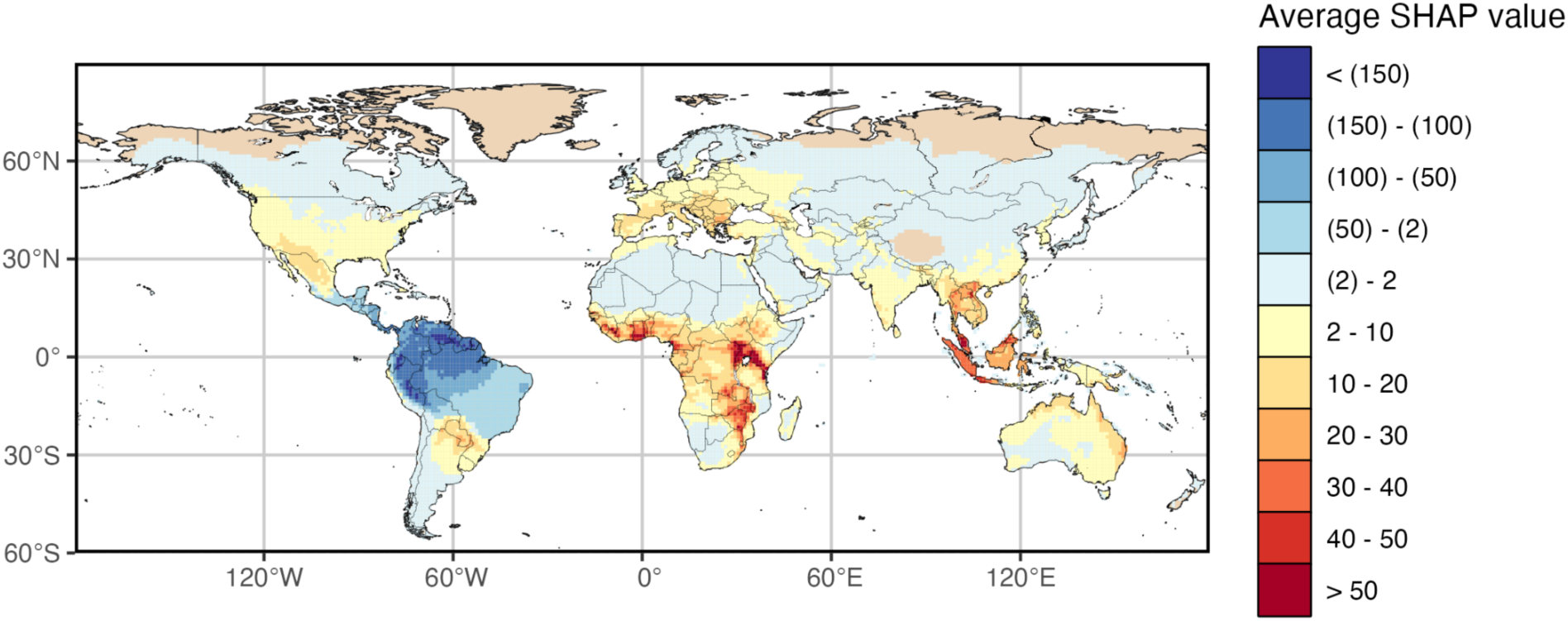
A map showing the geometric mean of average SHAP values for our ebolavirus (EBOV) model results across all predicted species. These SHAP values take into account all variables used in training, including phylogenetic, morphological, geographic, reproductive, and diet characteristics. Bluer pixels show more negative SHAP values (representing decreased binding strength), yellow pixels represent neutral values, and red pixels show positive SHAP values (representing increased binding strength). We used the geometric mean to reduce outliers caused by areas with little bat diversity.

Model results and exploratory figures and tables for each of the models described here can also be viewed interactively in greater detail in a Markdown output file available in a FigShare collection compiling model results (https://figshare.com/s/911bfcb78d49757902f9). Additionally, all scripts related to this study can be found on our Github repository (https://github.com/HanLabDiseaseEcology/bat-npc1-filovirus-modeling/).

## DISCUSSION

Predicting zoonotic risk depends on accurately identifying which host species are biologically susceptible to infection, yet susceptibility is rarely predicted using frameworks that integrate molecular mechanisms with macroecology. By combining experimentally grounded measures of filovirus-NPC1 binding strength with trait-based and phylogenetically informed modeling, we generated virus-specific predictions of binding strength across the global diversity of bats. Across models, predicted binding strength showed strong phylogenetic structuring alongside consistent contributions from life-history and ecological traits, indicating that receptor-mediated host-virus compatibility is evolutionarily conserved but modulated by species’ functional characteristics. Importantly, model predictions aligned with limited available binding assay data while extending inference to unsampled species and regions, revealing widespread and sometimes unexpected susceptibility across multiple bat lineages. Taken together, this framework advances a molecular biogeography of host susceptibility, linking receptor-level viral entry mechanisms to global patterns of bat host distribution.

### Phylogenetic structure and biogeographic context of susceptibility

The largest driver of predictive variation across all models was a suite of common phylogenetic variables, with the highest binding strengths concentrated in Molossidae (51.7% of species in the 90th percentile), followed by Pteropodidae (17.6%), Vespertilionidae (12.3%), and Hipposideridae (6.1%). This strong phylogenetic structuring is consistent with previous work documenting clade-level patterns in filovirus susceptibility [5,44] and earlier predictive efforts that similarly highlighted species within these families [4]. However, these studies necessarily integrated heterogeneous datasets including serology, PCR detections, and experimental infections, which vary substantially in confidence and comparability across filoviruses. By contrast, the use of receptor binding information provides a more mechanistically-grounded lens on susceptibility, and refines expectations for groups such as vespertilionid bats, which have historically shown limited surveillance evidence despite emerging as potentially susceptible in our predictions.

The clustering of confirmed and predicted high binding strength species within particular bat clades suggests that filovirus susceptibility reflects a long evolutionary history shaped by both host diversification and historical biogeographic processes. Although filoviruses were once thought to be evolutionarily recent [45,46], analysis incorporating purifying selection and the discovery of endogenous genomic elements indicates a much older origin, consistent with bat diversification across Miocene and Plio-Pleistocene biogeographic transitions [47,48].

Within bats, Asian and African faunal exchanges commonly occurred during the Miocene, providing repeated opportunities for dispersal and diversification across regions that now overlap with areas of documented filovirus activity. The expansion of Old World paleo-savanna biomes likely facilitated movement of more arid-adapted species across broad geographic areas [49], while intermittent humid periods across the Middle East and North Africa enabled additional crossings into Africa, including by pteropodid lineages [50,51]. These movements were modulated by species’ traits, with larger-bodied bats better able to traverse major barriers such as large rivers and arid regions, and cave-roosting bats able to move faster because they were less dependent on the expansion of tree species needed to provide roosting sites [50,52]. Subsequent Plio-Pleistocene forest expansion and contraction further isolated populations within refugial forest blocks, particularly in West Africa, promoting speciation patterns that remain detectable in contemporary bat phylogenies [52,53]. Similar refugial and dispersal-driven diversification dynamics have also been documented in other highly mobile taxa, including birds, indicating that these macroevolutionary responses to climatic oscillations are not unique to bats [31,32].

Importantly, these historical biogeographic processes overlap geographically with present-day hotspots of filovirus detection and provide a plausible macroevolutionary context for the strong phylogenetic structuring of binding strength observed in our models. In this sense, the widespread susceptibility patterns we predict across pteropodid and molossid lineages may reflect legacy effects of dispersal, diversification, and ecological filtering rather than solely contemporary surveillance biases. Consistent with this interpretation, modern analyses linking EBOV outbreaks to forest fragmentation [29] and accelerated substitution rates for EBOV and SUDV driven by land use change [28] suggest that contemporary anthropogenic change may be interacting with deeper refugial and biogeographic histories to shape the current zoonotic landscape.

### Functional trait correlates of receptor-mediated susceptibility

Our results also reveal a trait profile of binding compatibility between bat hosts and filoviruses that is grounded in host morphology and geography. However, in contrast to relatively consistent phylogenetic signal in most of our models, these trait-based patterns show greater variation in the importance and direction of individual trait effects on binding strength across filovirus models. The most consistent shared signals across models are geographic and morphological: higher predicted binding strength is typically associated with more arid environments and lowland or lower elevation regions, while lower relative wing loading also emerges as a broadly recurring characteristic. This differs from past modeling studies that have identified life history pace and diet as important predictors of reservoir status [4,5], though these results are not directly comparable, as our models use receptor binding strength rather than binary host or reservoir status as the response variable.

At first glance, the association between higher predicted binding strength, more arid conditions, and lower relative wing loading appears somewhat contradictory, because lower relative wing loading is typically associated with flight maneuverability of bats living in denser habitats, whereas arid environments often have a more open habitat structure. However, the aridity values in our models were calculated based on the year of occurrence (extracted from occurrence-based year-specific climate layers), which means they may be capturing relatively transient climatic conditions experienced by species rather than longer-term habitat characteristics. This interpretation is consistent with previous work showing that arid periods within wetter regions are associated with increased EBOV spillover risk [24]. Meanwhile, relative wing loading is linked to habitat use, maneuverability, and dispersal capacity in bats, which reflects cumulative rather than transient features of species’ environmental adaptations [54].

Given a potentially deep evolutionary history of filoviruses and the strong phylogenetic signal in our models, the recurring geographic profile across viruses may reflect bat lineages that diverged in part through refugial dynamics. Work on recent EBOV evolution supports the role of land use change in driving diversification of EBOV strains and suggests that ecological disruption may have enabled escape, and possible host switching, from a more spatially-restricted refugial area [28]. In contrast, other studies integrating latent infection support an older timeline of EBOV diversification and instead implicate reservoirs that use dense but geographically dispersed roosting networks [55]. Although this ecological profile appears superficially consistent with species such as *Rousettus aegyptiacus* and *Eidolon helvum*, experimental challenge studies show little evidence for onward transmission in *R. aegyptiacus*, and *in vitro* analyses suggest *E. helvum* may be refractory to EBOV infection [19,56,57], thereby implicating other widespread species as more plausible candidates.

An additional, speculative possibility is that the aridity signal we observed reflects broader physiological differences associated with NPC1-mediated cholesterol homeostasis. NPC1 is central to intracellular cholesterol trafficking [58], and cholesterol strongly shapes physical properties and fusion behavior of animal cell membranes [59]. Notably, direct cholesterol interaction has also been reported for EBOV GP2 during membrane fusion [60]. But more generally, membrane lipid composition is often tightly regulated to preserve function under environmental stress [61]. Thus, the variation we observed in predicted binding strength across bats may therefore partly capture differences in membrane homeostasis associated with their persistence in environments with dynamic moisture and temperature regimes, although this idea remains to be tested directly.

### Global patterns of predicted susceptibility and surveillance implications

Species predicted to have high binding strength are globally distributed across bat taxa and geographic regions, rather than being restricted to historically recognized filovirus outbreak zones (Figure 5). This broad pattern is consistent with previous predictive efforts [4] and suggests that the broad geographic signal is not merely an artifact of pooling data or modeling at the viral family level, but may instead reflect more broadly distributed potential for filovirus-host compatibility across diverse bat lineages.

Filoviruses are not confined to where outbreaks have been documented, and mounting evidence indicates a much broader geographic and evolutionary footprint, including discoveries in Europe and Asia [21,22,62], as well as filovirus-like genomic elements detected in a wider range of bat taxa such as microbats [63], suggesting a history of past exposure that led to the integration of filoviral sequences into host genomes. Moreover, paleoviral analyses indicate that the inclusion of these filovirus-like elements have not been restricted to bats, with similar genomic signatures identified in rodents and marsupials [64,65].

Evidence from recent virological studies further underscores the geographic and taxonomic breadth of filoviruses. Among vertebrates, fish filoviruses have been found in Antarctica [66], Switzerland [67,68], China [69], Germany [70], and Australia [71], and a snake-associated filovirus was discovered in a *Bothrops atrox* in Brazil [72]. Within mammals, filovirus sequence fragments have been detected in *Nyctinomops laticaudatus* in Brazil [73] alongside serologic evidence for filovirus exposure in bats from Trinidad and Tobago [74]. Taken together, these findings suggest that filoviruses and filovirus-like elements extend beyond geographic regions and hosts represented by outbreak records alone.

### Data limitations and sources of discordant predictions

Although these models represent a step forward in predicting filovirus susceptibility across bats, their performance is still constrained by persistent data deficiencies. In particular, models for MLAV and RESTV performed poorly across multiple evaluation metrics including both RMSE and R^2^. For SUDV and TAFV, correlation-based metrics were similarly lacking despite stronger RMSE values, suggesting that while absolute prediction error may be moderate, the models struggle to reliably rank species relative to one another. This likely reflects a clumped distribution of binding strengths for these viruses, which limits the ability of correlation metrics to distinguish fine-scale differences among species.

More broadly, small sample sizes and uneven taxonomic representation influence model behavior, especially for groups that are poorly represented or absent in the training dataset, such as Miniopteridae and the greater diversity within Hipposideridae. There are no miniopterids with phylogenetic information in our training dataset, and the most informative phylogenetic PCo axes primarily differentiate species within the genus *Hipposideros* rather than capturing phylogenetic variation across Hipposideridae more broadly (Supplementary Figures 1-2). These limitations identify where additional empirical data are needed to stabilize estimates and refine predictions. With additional binding assay data or sequence-based predictions of binding strength, model performance for these viruses is expected to improve.

Additionally, it is possible that the trait data compiled here are insufficient to fully resolve the biological patterns underlying host binding strength. As currently constructed, most trait databases report species-level averages, which can smooth over seasonal and geographic variability and obscure within-species differences, particularly for recently described or taxonomically complex groups [75,76]. Miniopterids, which produced discordant predictions in our analyses, illustrate this problem clearly. Binding assay data for this family come from a newly described species with limited trait information and no representation in published mammalian phylogenies. Consequently, the models lacked training data for the most important predictors. In the absence of trait and phylogenetic variables, the algorithms inferred miniopterid susceptibility indirectly through their position on phylogenetic PCo axes, which also capture high binding strength in other bat families. This indirect inference likely generated an apparent signal of high susceptibility for miniopterids that reflects extrapolation from incomplete trait and phylogenetic data, rather than evidence specific to miniopterids.

### Interpretation of species-level predictions

Our modeling approach captures broad patterns in binding strength that are anchored in experimentally-derived binding assay values [37]. Compared to earlier efforts to predict filovirus susceptibility, these trait-based models provide finer resolution at the viral species level rather than relying on pooled family-level signals. For most of the filoviruses modeled here, we accurately predict binding strength (Tables 1-2, Supplementary Figure 3), with the strongest performance observed in the *Orthomarburgvirus*-like clade. Some of this strong performance may reflect features of the training data, including the relatively high representation of Phyllostomidae (37.5%) and the use of physicochemical predicted binding strength values as labels for several viruses, which may make those models more likely to recover labels more closely tied to underlying sequence-based predictions. In contrast, our model of LLOV, which combined experimentally observed and predicted labels, was less well explained by functional trait data. This difference suggests that trait-based predictability may vary among filovirus clades and that binding strength in the *Orthoebolavirus-*like clade may be shaped by a more complex or less easily captured combination of ecological and evolutionary factors. Even where these models primarily recover binding strength patterns derived from physicochemical properties, they still provide a useful bridge that extends susceptibility inference to species lacking sequence or binding assay data.

Importantly, model outputs also show strong biological concordance, with many species previously associated with specific filoviruses ranking among the higher predicted binding strength values. For example, the known reservoir for DEHV, *Rousettus leschenaultii* [22], falls within the 80th percentile of predicted binding strength results (S7 Table). Although the EBOV reservoir is unknown, *Mops condylurus* in challenge studies show viral replication and shedding and rank in the 99th percentile of predictions here [77]. For MARV, *Rousettus aegyptiacus* and *Miniopterus inflatus* both rank in the 80th percentile (note that *Rhinolophus eloquens*, however, is in the 40th percentile). Of these, *R. aegyptiacus* is an established MARV reservoir while the latter two species have been reported as PCR-positive [78,79]. The bats associated with LLOV (*Miniopterus schreibersii* and *Miniopterus pusillus*) and BOMV (*Mops condylurus and Chaerephon pumilus*) consistently fall in the upper ranges of predicted binding strength (70th-98th percentiles) [20,21,41].

Predicted binding strength reflects receptor compatibility rather than direct zoonotic transmission risk or reservoir status. We interpret higher binding strength as indicating greater potential susceptibility rather than confirmed zoonotic capacity because binding between NPC1 and filovirus glycoprotein represents only the entry stage of infection and does not account for downstream processes such as replication, shedding, or immune modulation that also determine reservoir competence [80–83]. Nevertheless, because it is a necessary first step to viral entry, receptor binding provides a mechanistically informed and scalable starting point for comparing susceptibility across diverse host-pathogen systems beyond filoviruses [84–86].

### Cross-virus similarity in predicted susceptibility

Binding strength predictions are highly correlated across many filoviruses, especially among the *Orthomarburgvirus*-like filoviruses (DEHV, MARV, and RAVV), suggesting that cross-virus similarity reflects shared structure in the training data as well as underlying biological relatedness. This interpretation is consistent with viral phylogeny. Although there are several filovirus genera associated with mammals (*Cuevavirus*, *Dianlovirus*, *Orthoebolavirus*, *Orthomarburgvirus*), there are currently two main clades dominated by species from either *Orthoebolavirus* or *Orthomarburgvirus*. DEHV is currently a species in the genus *Dianlovirus*, but binding predictions show a close association with *Orthomarburgvirus* that match the closer relationship to this genus [87], whereas the lack of similarly high correlation to another *Dianlovirus*, MLAV, may reflect weaker model performance for this species. More broadly, similarities in species-level binding predictions across filoviruses are consistent with evidence for high serological cross-reactivity, particularly among *Orthomarburgvirus*-like filoviruses [22,34,88]. The similarity in binding strength across multiple filoviruses within the same host species suggests that some regions may contain bat assemblages with potential compatibility for several filoviruses at once, particularly in West and East Africa where multiple filoviruses have been documented in close proximity (Figure 6). In such settings, surveillance may benefit from multiplexed approaches that may be more informative than targeting single viruses.

**Figure 6.**
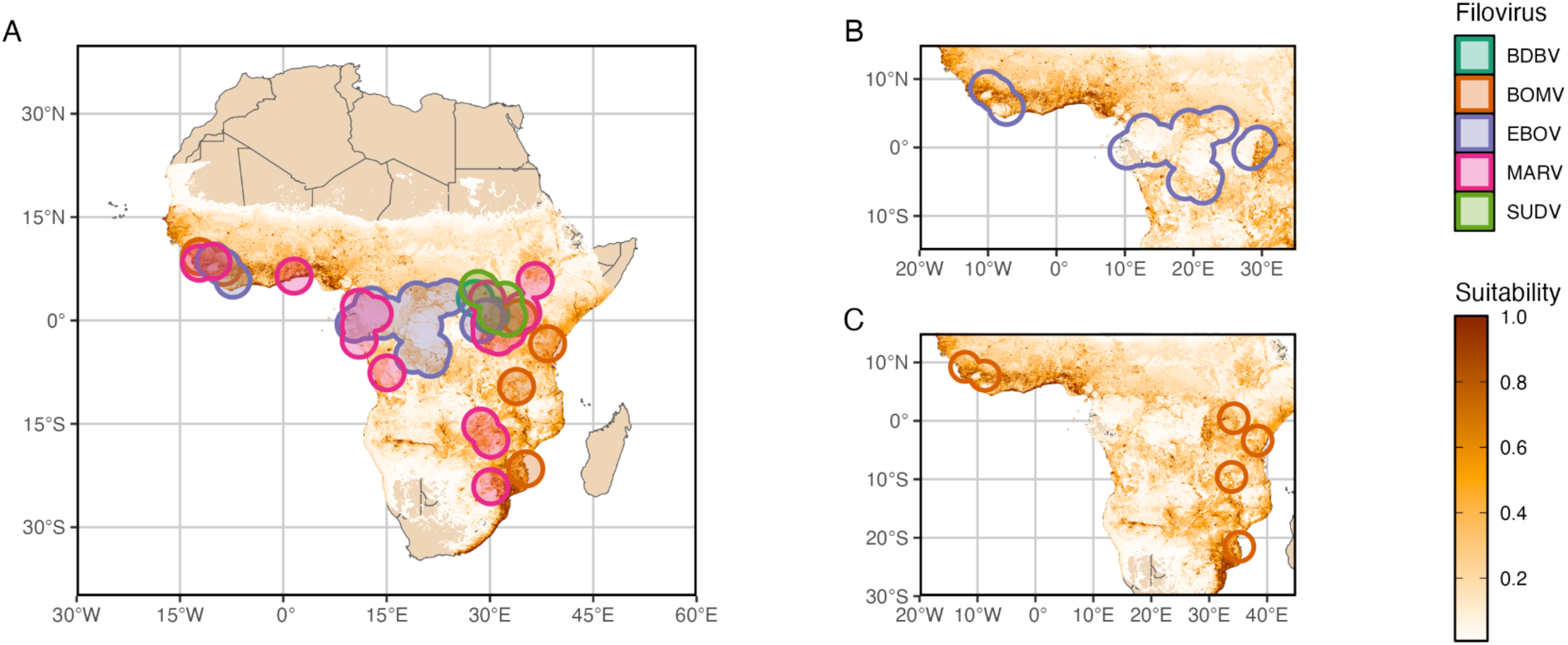
A map of Africa showing an example of one species with high binding strength to multiple filoviruses, *Mops condylurus*, and all known African filovirus occurrences. These occurrences, which are either outbreaks or surveillance records from live animals, are buffered by 50 km to account for animal and human movement. In all panels, habitat suitability for *M. condylurus* is shown in darker orange colors, estimated from an ecological niche model at 5 km resolution based on occurrences from the African Bat Database <u>(Monadjem et al. 2024)</u>. Circles in panels B and C show Ebola and Bombali virus occurrences, respectively, and identify areas that do (darker orange) and do not (white) appear to coincide with the geographic range distribution for *M. condylurus*.

## Conclusions

These predictions are best interpreted as approximating the fundamental host range of filoviruses, based on NPC1-mediated binding associated with viral entry rather than the full suite of processes required for successful infection, transmission, and maintenance in reservoir populations. Some of the patterns identified here may also reflect historical biogeography and evolutionary contingency, without implying current associations. By contrast, realized host range will depend on additional cellular, ecological, and epidemiological constraints, as well as spatial overlap between viruses and their potential hosts. Identifying realized reservoirs therefore remains a major challenge that will benefit from an iterative, model-guided framework that integrates surveillance, experimental assays, ecological data, and comparative modeling, thereby completing the loop between field data, hypothesis generation, prediction, and empirical validation as inference improves over time [89].

Ongoing advances in metagenomic and targeted sequencing, coupled with expanded surveillance across understudied regions and taxa, will be important for refining these predictions and resolving data gaps highlighted here [90]. More broadly, the widespread and phylogenetically structured susceptibility patterns we identify suggest that evolutionary history and biogeographic context are important for assessing zoonotic risk. In an era of rapid environmental change driven by land use shifts, climate change, and global species movement, a clearer understanding of host evolutionary and biogeographic history may provide an important foundation to better anticipate how the host range and distribution of filoviruses, and other emerging viruses, could shift in the future.

## METHODS

To determine trait profiles indicating susceptibility to bat-borne filoviruses and to predict NPC1 binding strength across bat species, we followed a workflow building on previous comparative modeling approaches [8,36]. Our analytical pipeline proceeded in four stages: (i) compilation and harmonization of ecological, phylogenetic, and environmental trait data for bats, (ii) integration of experimentally derived and predicted NPC1 binding strength labels, (iii) boosted regression modeling of virus-specific binding strength using trait and phylogenetic predictors, and (iv) extrapolation of predictions across all bat species and geographic contexts (Figure 2). Because the random forest model of binding strength in Lasso et al. [37] was trained almost exclusively on bats, we restricted our analyses to Chiroptera to ensure consistency between label structure and predictor space.

### Trait data

We compiled mammalian functional trait data using the COMBINE trait dataset as the primary backbone [76]. COMBINE harmonizes multiple established mammalian trait databases (e.g., PanTHERIA [75], EltonTraits [91]) and is better aligned with recent taxonomic updates and additions [92]. To maximize trait coverage across bat species, we used the imputed version of the dataset and one-hot encoded categorical variables for compatibility with downstream modeling (S1 Appendix).

Given the unique functional space that bats occupy among mammals and a robust literature on bat-specific traits, we supplemented COMBINE with variables or additional coverage from other studies [2,93,94], crucially introducing several morphological and roosting variables. Habitat association data were obtained from the IUCN Red List ([95]; version 2023-1) using the rredlist package (version 0.7.1) in R version 4.3.2 [96,97], capturing suitability across broad habitat classifications (e.g., forest, savanna, shrubland, etc.) as well as more specific associations with moist forest and moist savanna categories relevant to hypotheses linking aridity and seasonal rainfall to EBOV spillover [24].

To further investigate the role of aridity, we downloaded all georeferenced bat occurrence records from the Global Biodiversity Information Facility (GBIF) for the years 1958-2023 [98]. These records were filtered for identified geographic issues and for an uncertainty radius under 5 km. Using the dplyr, sf, and terra packages [99–102], we extracted the aridity index for each occurrence record and summarized these per species as the minimum aridity index found at an occurrence point, the mean aridity index across all occurrence points, and the threshold that includes 80% of all aridity index values. We used monthly precipitation and potential evapotranspiration data from Terra Climate to calculate the yearly aridity index for 1958-2023 [103].

We incorporated phylogenetic information by taking the first 17 axes of a phylogenetic principal coordinates analysis (PCoA). Briefly, we used a random subset of 1,000 phylogenetic trees from Upham et al. ([104]; Figure 2). For the broadest coverage in our dataset, we used the set of trees which imputed positions of species without sequence information, necessitating the use of a large subset of trees. We calculated the pairwise phylogenetic distances between species by creating a distance matrix for each tree using branch lengths with the ape package in R (version 5.7.1; [105]). We took the mean distance values across all 1,000 trees to create a consensus distance matrix representing the average pairwise phylogenetic distances across all trees. We then subset this distance matrix to bats and performed a phylogenetic principal coordinates analysis (PCoA) on this consensus matrix to reduce the dimensionality of the data to 17 axes comprising 90% of the variation using the ape package. Although harder to interpret than binary variables indicating taxonomic family, initial axes, such as PCo1 and PCo2, typically describe broader relationships within bats (e.g., Yinpterochiroptera and Yangochiroptera, family boundaries) and later axes (e.g., PCo17) separate genera or species within these families. To address taxonomic mismatches throughout data collation, we consulted the Mammal Diversity Database [92].

Before model parameterization and evaluation, we checked variables for high levels of skewness, near zero variation, and high degrees of missingness. We used the moments (version 0.14.1; [106]) package to calculate skewness. For heavily skewed variables, defined as an absolute skewed value of 1.5, we applied log transformation. We removed all variables with near zero variation using the caret package (version 6.0-94; [107,108]) along with all variables that are missing data in over 30% of the records.

### Boosted regression

We used XGBoost, a gradient boosting method (xgboost R package, version 1.7.6.1; [109]), to model virus-specific binding strength as a function of species-level ecological and phylogenetic traits. As filoviruses span a large geographic area and observed binding strength values for a bat could differ widely across viruses, we trained separate models for each filovirus. The training dataset was restricted to bat species with experimentally derived NPC1 binding strength estimates from Lasso et al. [37] (81 species, reduced to 72 after accounting for taxonomic mismatches and unavailable trait data). Note that Lasso et al. [37] used a random forest model to predict binding strength values for Dehong virus (DEHV), Marburg virus (MARV), and Ravn virus (RAVV), so all labels used for training models of these viruses were predicted rather than experimental. A separate set of 54 records with binding strength values derived from the random forest model trained on physiochemical properties by Lasso et al. [37] was retained as a fully held out validation dataset.

Given limited sample size, all parameterization, evaluation, and model interpretation steps were done using 100 bootstrap iterations. Within each iteration, data were partitioned into random subsets of training (75% of the data, 54 rows) and testing data (25% of the data, 18 rows), while preserving the distribution of binding strength values. We used a grid search method to parameterize the eta, interaction depth, minimum child weight, subsampling ratio, columns selected per tree, gamma, and number of trees in XGBoost (Table 1). Trees were bounded to a minimum of 1,000 and maximum of 10,000 and the eta was similarly bounded to a minimum of 0.001 and maximum of 0.5. We used a Latin hypercube sampling approach to select 500 combinations of these parameters to evaluate and choose from.

Model performance was evaluated using four-fold cross validation implemented in the tidymodels (version 1.1.1) and parsnip (version 1.1.1) packages in R [110,111], with parameter selection based on minimizing root mean square error (RMSE) and maximizing R^2^. After selecting optimal hyperparameters for each filovirus-specific model, final models were fit to the full training dataset and evaluated on the held out validation dataset (Table 2).

For model interpretation of each bootstrap subset, we calculated Shapley values for each predictor using the SHAPforxgboost package (version 0.1.3; [112]) in R. Models trained on the full training dataset were used to predict binding strength across 1,342 bat species with available trait data.

### Outbreak and virus detection locations

We compiled outbreak and virus detection locations by building on the dataset of EBOV occurrences assembled by [24], and extending it to 2025 using CDC and WHO reporting on confirmed outbreaks of filoviruses. Because many filoviruses are not currently known to cause human outbreaks, we supplemented outbreak records with virus detections in animals using targeted searches. To ensure taxonomic specificity, we restricted detections to those supported by virus isolation or PCR evidence, excluding serology results due to known cross reactivity among filoviruses [34,113]. For mammal distribution data, we used polygons from [114]. We used the rnaturalearth package for administrative boundary data [115].

## ACKNOWLEDGMENTS

The authors thank Andrew Kramer for several helpful comments that increased the readability of this manuscript. AAC and BAH are funded by the joint NIFA-NSF-NIH Ecology and Evolution of Infectious Disease award 2023-70432-40381.

## SUPPORTING INFORMATION

**S1 Appendix.** A table of all variables used for our trait-based modeling of binding strength. For each variable, we give the variable name, the units the variable is calculated in, the percent coverage across our training dataset, whether the variable was retained, whether the variable was log transformed due to a skewed distribution, and the source of the variable. We retained the variable names from their sources to allow for easy followup with the original source for detailed descriptions or calculations.

**S2 Figure.**
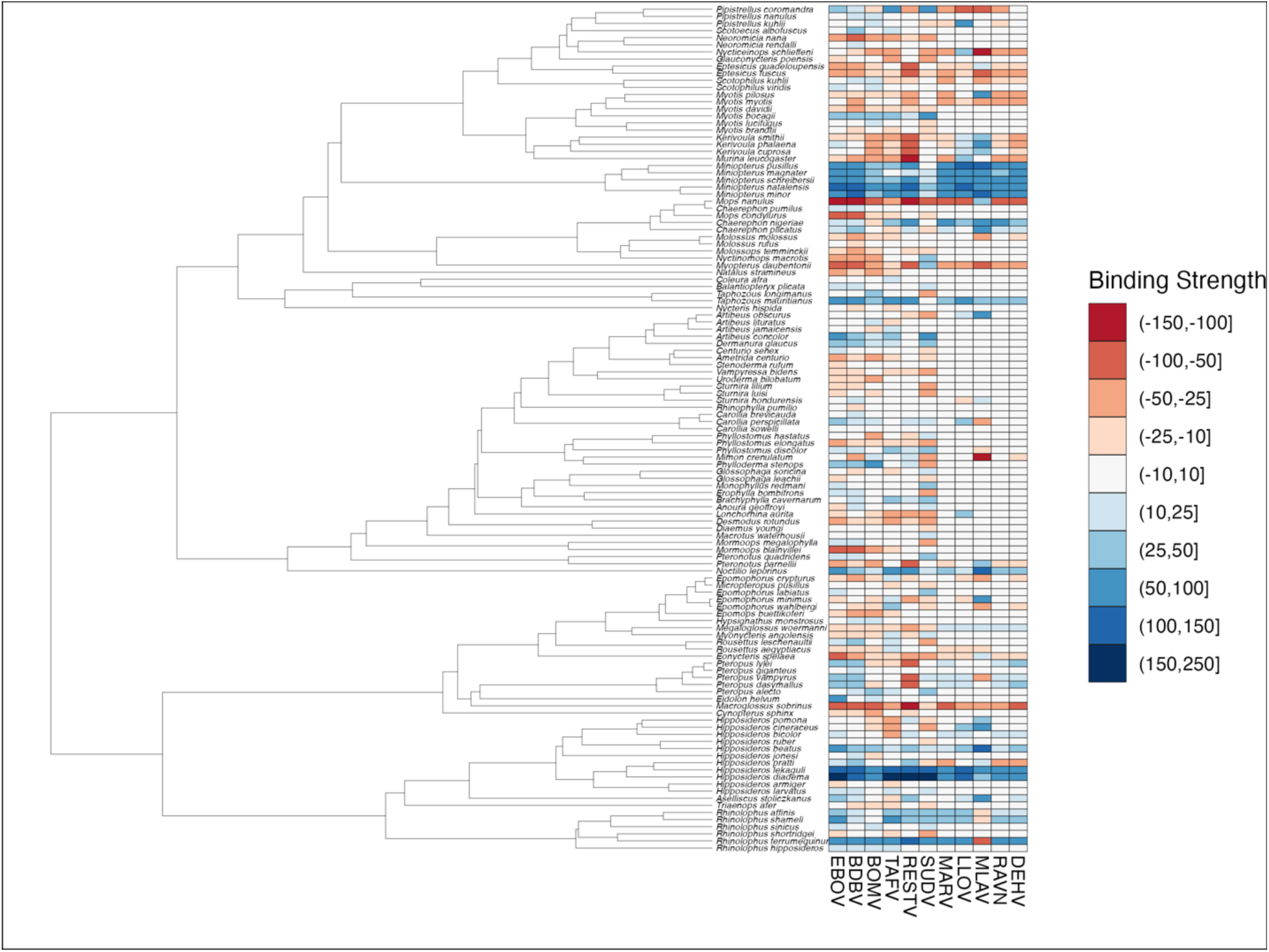
A phylogenetic tree of all species in our training and validation datasets with a heatmap showing the difference between observed and predicted binding strength for 11 filoviruses. The phylogenetic tree is from Upham et al. 2019. In the heatmap, redder shades depict a lower predicted binding strength than the observed and bluer shades show a higher predicted binding strength than observed.

**S3 Figure.**
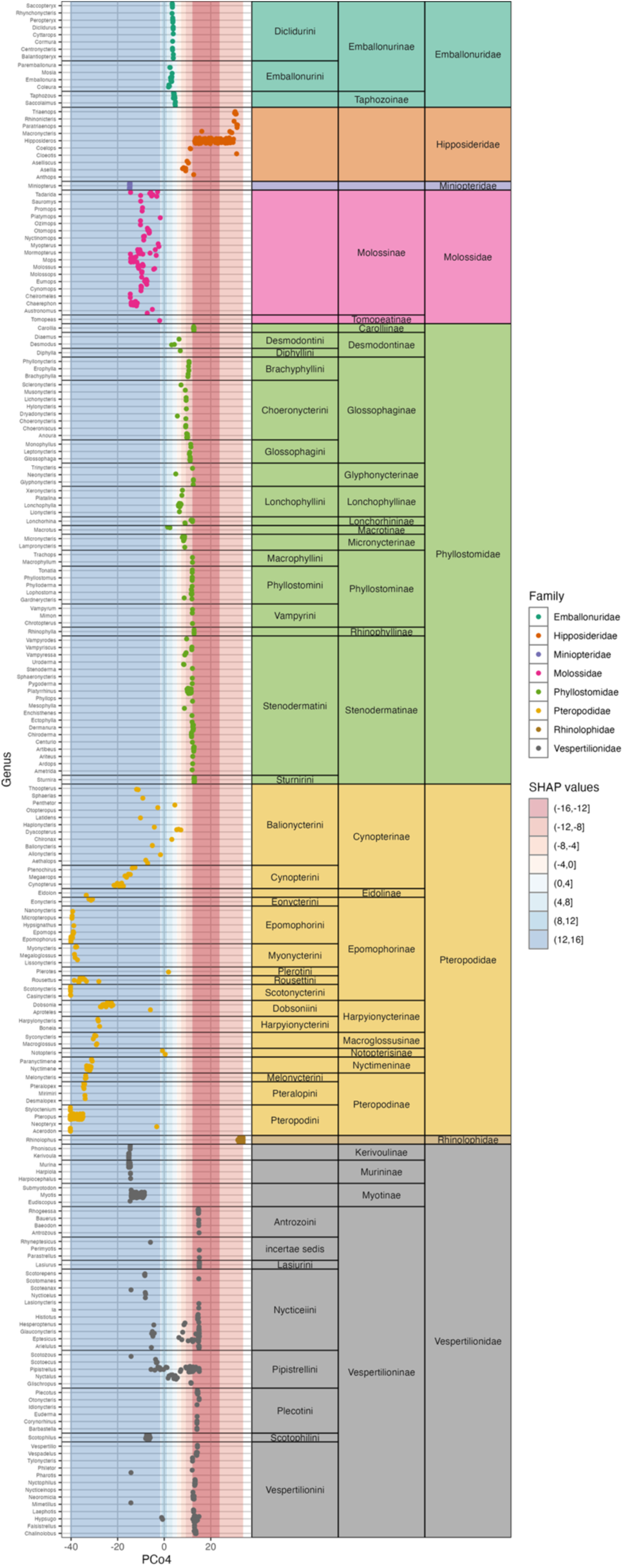
Panels in this plot show the distribution of PCo4 axis values (x-axis) for species in a particular genus (y-axis). Bat families are color coded and genera are arranged by family, subfamily, and tribe as seen in the rightmost panels. The shading under the points shows the SHAP values associated with those PCo4 axis values, with redder values relating to axis values associated with a decrease in binding strength and bluer values relating to axis values associated with an increase in binding strength.

**S4 Figure.**
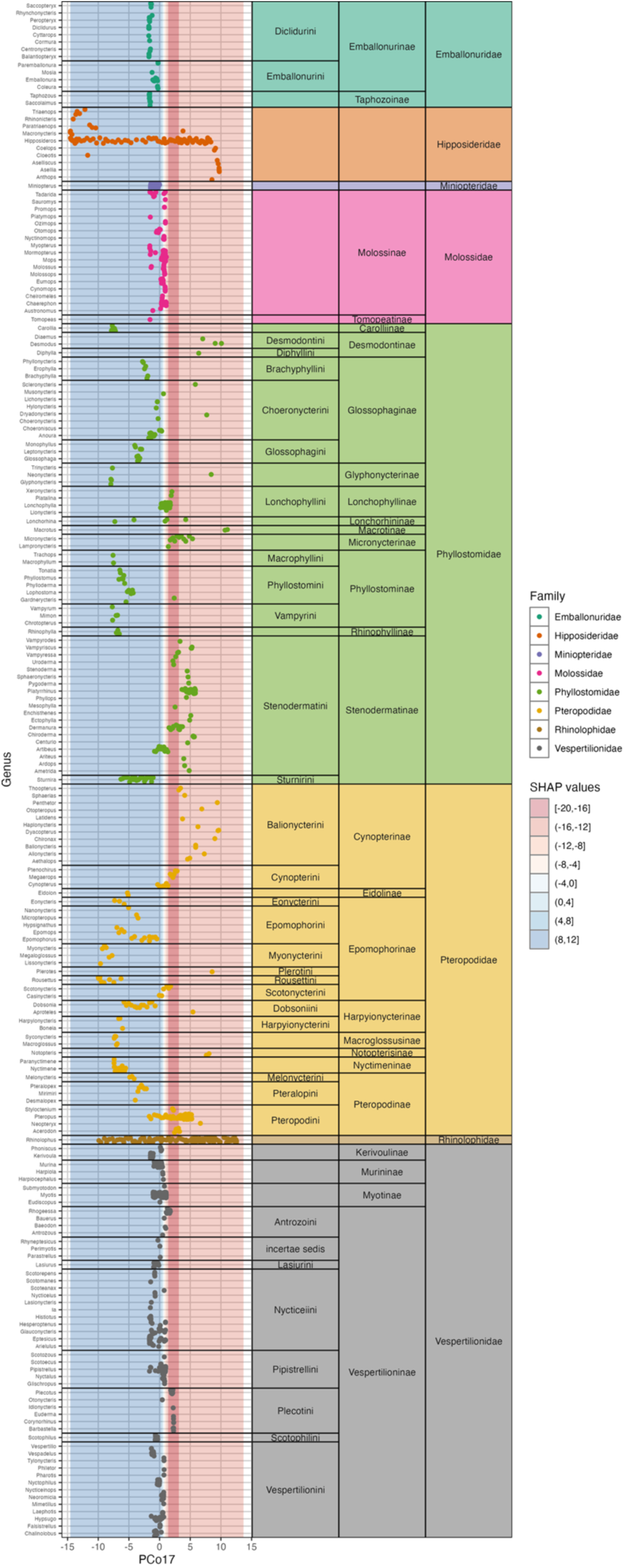
Panels in this plot show the distribution of PCo17 axis values (x-axis) for species in a particular genus (y-axis). Bat families are color coded and genera are arranged by family, subfamily, and tribe as seen in the rightmost panels. The shading under the points shows the SHAP values associated with those PCo17 axis values, with redder values relating to axis values associated with a decrease in binding strength and bluer values relating to axis values associated with an increase in binding strength.

**S5 Figure.**
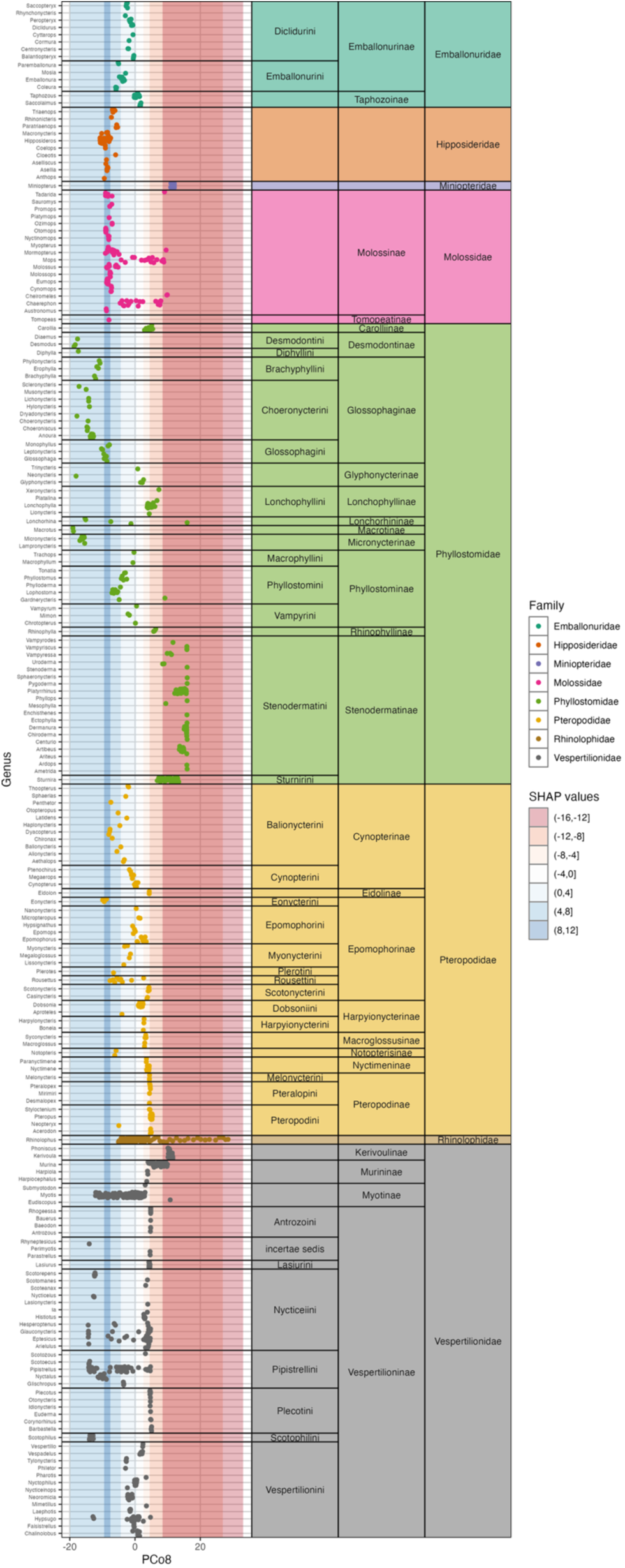
Panels in this plot show the distribution of PCo8 axis values (x-axis) for species in a particular genus (y-axis). Bat families are color coded and genera are arranged by family, subfamily, and tribe as seen in the rightmost panels. The shading under the points shows the SHAP values associated with those PCo8 axis values, with redder values relating to axis values associated with a decrease in binding strength and bluer values relating to axis values associated with an increase in binding strength.

**S6 Figure.**
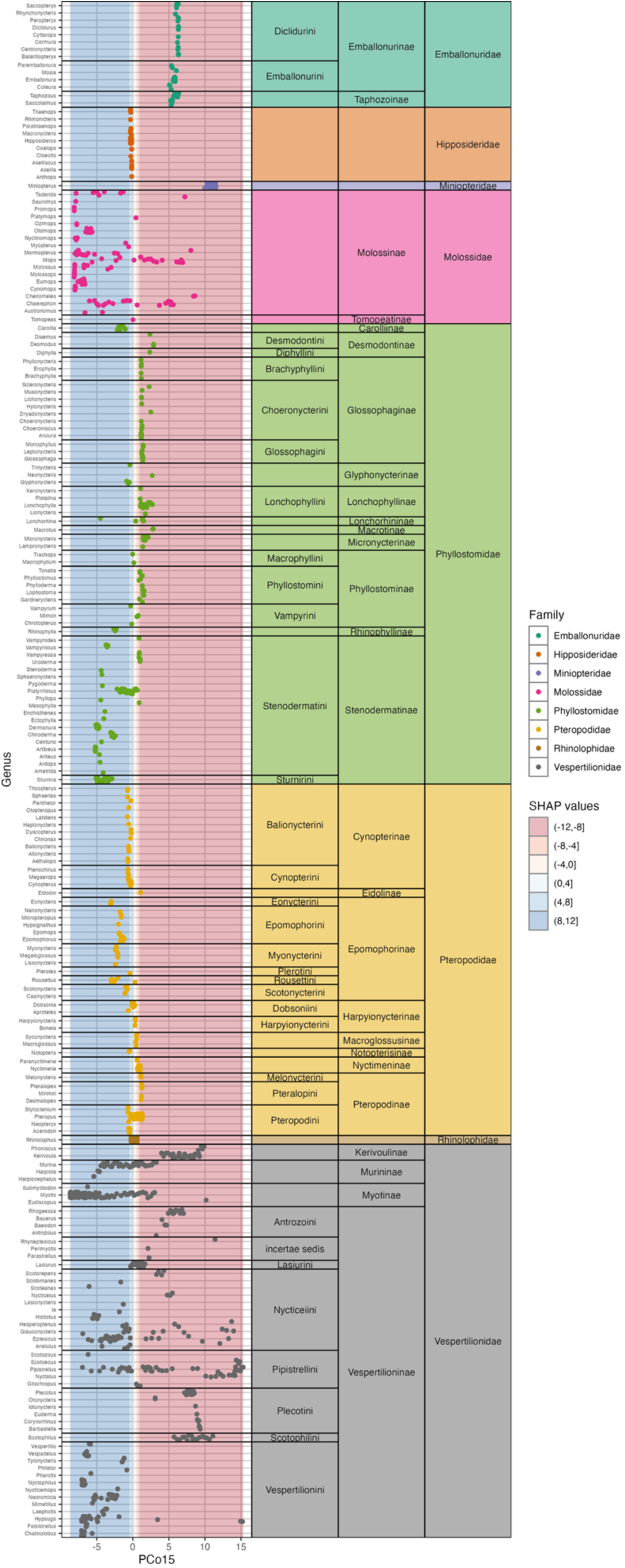
Panels in this plot show the distribution of PCo15 axis values (x-axis) for species in a particular genus (y-axis). Bat families are color coded and genera are arranged by family, subfamily, and tribe as seen in the rightmost panels. The shading under the points shows the SHAP values associated with those PCo15 axis values, with redder values relating to axis values associated with a decrease in binding strength and bluer values relating to axis values associated with an increase in binding strength.

**S8 Figure.**
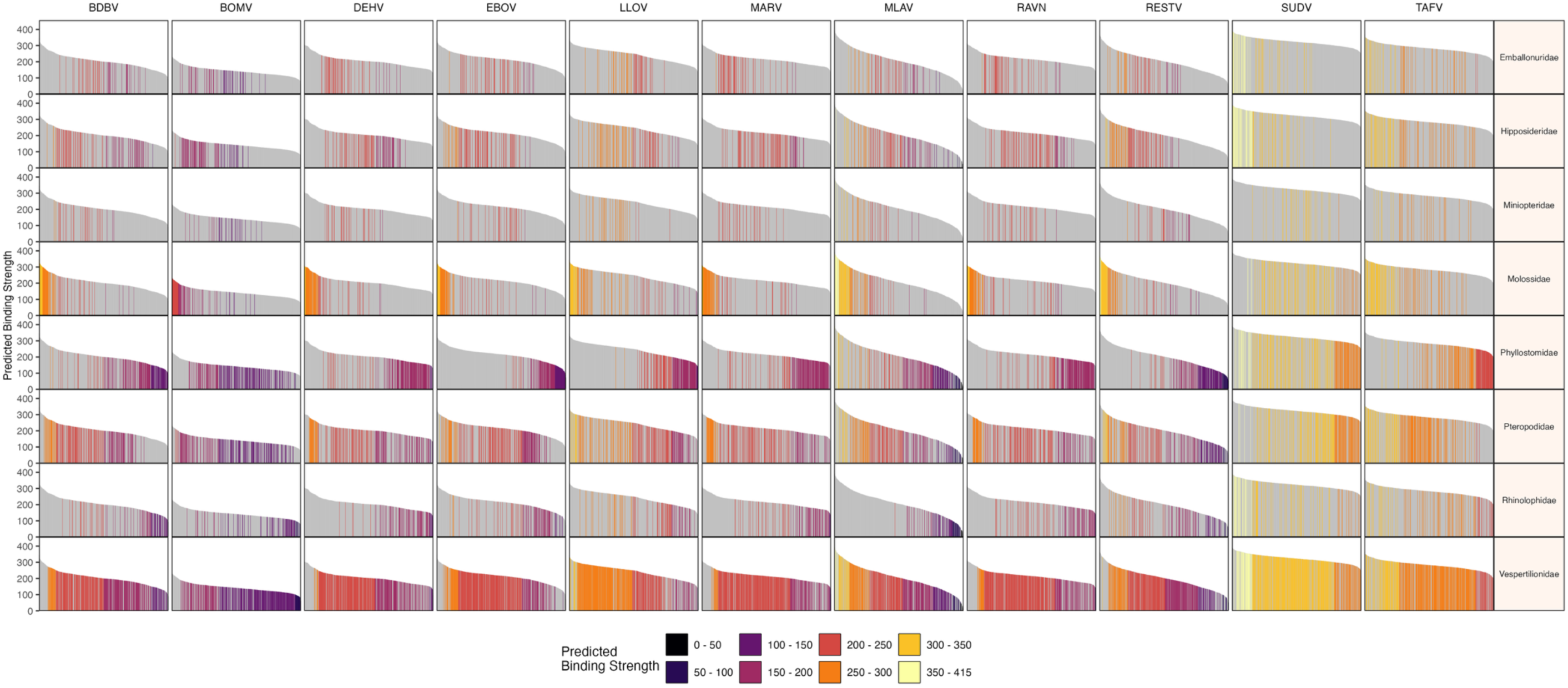
Model predictions presented for our 11 filoviruses (columns) across all bats shown for the eight most speciose bat families (rows). The y-axis shows predicted binding strength to EBOV for each bat NPC1, while the x-axis arranges the bat species in order from highest binding to lowest binding. Note that this order is necessarily different for each column. Each panel depicting a bat family shows only those species in the family with the color of the bar relating to a binned binding strength category.

**S7 Table.** A spreadsheet showing binding strength predictions for 1,342 bats for 11 filoviruses. Each row shows the predictions for a single bat species, with columns for its scientific name, family, and the 11 filovirus predictions.

