## Supplementary material for "From receptor binding to biogeography: Multi-scale prediction of filovirus hosts in bats": S1 Appendix

### **S1 Appendix - Variables**

A table of all variables used for our trait-based modeling of binding strength. For each variable, we give the variable name, the units the variable is calculated in, the percent coverage across our training dataset, whether the variable was retained, whether the variable was log transformed due to a skewed distribution, and the source of the variable. We retained the variable names from their sources to allow for easy followup with the original source for detailed descriptions or calculations.

| Variable | Units | Coverage (%) | Retained | Log transformed | Source |
| --- | --- | --- | --- | --- | --- |
| adult_mass_g | g | 100 | Y | Y | COMBINE |
| brain_mass_g | g | 98.6 | Y | Y | COMBINE |
| adult_body_length_mm | mm | 100 | Y | Y | COMBINE |
| adult_forearm_length_mm | mm | 91.8 | Y | Y | COMBINE |
| max_longevity_d | days | 98.6 | Y | N | COMBINE |
| maturity_d | days | 35.6 | Y | N | COMBINE |
| female_maturity_d | days | 98.6 | Y | N | COMBINE |
| female_relative_maturity | - | 98.6 | Y | Y | Han et al. 2016 |
| male_maturity_d | days | 19.2 | N | N | COMBINE |
| age_first_reproduction_d | days | 98.6 | Y | N | COMBINE |
| relative_age_first_birth | - | 98.6 | Y | Y | Han et al. 2016 |
| gestation_length_d | days | 98.6 | Y | N | COMBINE |
| teat_number_n | number | 0 | N | N | COMBINE |
| litter_size_n | number | 100 | Y | Y | COMBINE |
| litters_per_year_n | number | 98.6 | Y | N | COMBINE |
| interbirth_interval_d | days | 98.6 | Y | N | COMBINE |
| neonate_mass_g | g | 46.6 | Y | Y | COMBINE |
| weaning_age_d | days | 98.6 | Y | Y | COMBINE |
| weaning_mass_g | g | 20.5 | N | N | COMBINE |
| generation_length_d | days | 98.6 | Y | N | COMBINE |
| mass_specific_production_rate | - | 46.6 | Y | N | Hamilton et al. 2011 |
| dispersal_km | km | 0 | N | N | COMBINE |
| density_n_km2 | individuals/km^2^ | 0 | N | N | COMBINE |
| hibernation_torpor | binary | 98.6 | Y | N | COMBINE,  Luis et al. 2013 |
| fossoriality | binary | 98.6 | N | N | COMBINE |
| home_range_km2 | km^2^ | 0 | N | N | COMBINE |
| social_group_n | number | 6.8 | N | N | COMBINE |
| dphy_invertebrate | percent | 100 | Y | N | COMBINE |
| dphy_vertebrate | percent | 100 | N | N | COMBINE |
| dphy_plant | percent | 100 | Y | N | COMBINE |
| det_inv | percent | 95.9 | Y | N | COMBINE |
| det_vend | percent | 95.9 | N | N | COMBINE |
| det_vect | percent | 95.9 | N | N | COMBINE |
| det_vfish | percent | 95.9 | N | N | COMBINE |
| det_vunk | percent | 95.9 | N | N | COMBINE |
| det_scav | percent | 95.9 | N | N | COMBINE |
| det_fruit | percent | 95.9 | Y | N | COMBINE |
| det_nect | percent | 95.9 | Y | N | COMBINE |
| det_seed | percent | 95.9 | N | N | COMBINE |
| det_plantother | percent | 95.9 | N | N | COMBINE |
| det_diet_breadth_n | number | 100 | Y | N | COMBINE |
| trophic_level | ordered | 100 | Y | N | COMBINE |
| activity_cycle | ordered | 100 | N | N | COMBINE |
| freshwater | binary | 100 | N | N | COMBINE |
| marine | binary | 100 | N | N | COMBINE |
| terrestrial_non.volant | binary | 100 | N | N | COMBINE |
| terrestrial_volant | binary | 100 | N | N | COMBINE |
| upper_elevation_m | m | 49.3 | N | N | COMBINE |
| lower_elevation_m | m | 46.6 | Y | Y | COMBINE |
| altitude_breadth_m | m | 32.9 | N | N | COMBINE |
| island_dwelling | binary | 76.7 | Y | N | COMBINE |
| dissected_by_mountains | binary | 75.3 | Y | N | COMBINE |
| glaciation | binary | 75.3 | N | N | COMBINE |
| habitat_breadth_n | number | 98.6 | Y | N | COMBINE |
| foraging_aerial | binary | 100 | Y | N | COMBINE |
| foraging_arboreal | binary | 100 | Y | N | COMBINE |
| foraging_ground | binary | 100 | N | N | COMBINE |
| foraging_marine | binary | 100 | N | N | COMBINE |
| foraging_scansorial | binary | 100 | N | N | COMBINE |
| island_mainland | binary | 98.6 | Y | N | COMBINE |
| island_marine | binary | 98.6 | N | N | COMBINE |
| island_landbridge | binary | 98.6 | N | N | COMBINE |
| island_endemic | binary | 98.6 | Y | N | COMBINE |
| bio_palearctic | binary | 100 | N | N | COMBINE |
| bio_afrotropical | binary | 100 | Y | N | COMBINE |
| bio_antarctic | binary | 100 | N | N | COMBINE |
| bio_australasian | binary | 100 | N | N | COMBINE |
| bio_indomalayan | binary | 100 | Y | N | COMBINE |
| bio_nearctic | binary | 100 | Y | N | COMBINE |
| bio_neotropical | binary | 100 | Y | N | COMBINE |
| bio_oceanian | binary | 100 | N | N | COMBINE |
| iucn_forest | binary | 100 | Y | N | IUCN Red List |
| iucn_savanna | binary | 100 | Y | N | IUCN Red List |
| iucn_shrubland | binary | 100 | Y | N | IUCN Red List |
| iucn_grassland | binary | 100 | Y | N | IUCN Red List |
| iucn_wetlands | binary | 100 | N | N | IUCN Red List |
| iucn_rocky | binary | 100 | Y | N | IUCN Red List |
| iucn_caves | binary | 100 | Y | N | IUCN Red List |
| iucn_desert | binary | 100 | N | N | IUCN Red List |
| iucn_marineneritic | binary | 100 | N | N | IUCN Red List |
| iucn_marineoceanic | binary | 100 | N | N | IUCN Red List |
| iucn_marinedeep | binary | 100 | N | N | IUCN Red List |
| iucn_marineintertidal | binary | 100 | N | N | IUCN Red List |
| iucn_marinecoastal | binary | 100 | N | N | IUCN Red List |
| iucn_artificialterr | binary | 100 | N | N | IUCN Red List |
| iucn_artificialaquatic | binary | 100 | N | N | IUCN Red List |
| iucn_introducedveg | binary | 100 | N | N | IUCN Red List |
| iucn_moistforest | binary | 100 | Y | N | IUCN Red List |
| iucn_moistsavanna | binary | 100 | Y | N | IUCN Red List |
| min_aridity_index | - | 97.3 | Y | N | This study |
| mean_aridity_index | - | 97.3 | Y | N | This study |
| perc80_aridity_index | - | 97.3 | Y | N | This study |
| relative_wing_loading | - | 68.5 | Y | N | Crane et al. 2022, Guy et al. 2020 |
| aspect_ratio | - | 67.1 | Y | N | Crane et al. 2022, Guy et al. 2020 |
| median_aggregation_size | number | 42.5 | Y | Y | Guy et al. 2020 |
| migration | binary | 34.2 | Y | N | Luis et al. 2013 |
| mixed_species_association_n | number | 26.0 | N | N | Guy et al. 2020 |
| cave_roost | binary | 69.9 | Y | N | Guy et al. 2020 |
| cavity_roost | binary | 69.9 | Y | N | Guy et al. 2020 |
| foliage_roost | binary | 69.9 | Y | N | Guy et al. 2020 |
| generalist_roost | binary | 69.9 | Y | N | Guy et al. 2020 |
| pco_1 | - | 98.6 | Y | N | Upham et al. 2019 |
| pco_2 | - | 98.6 | Y | N | Upham et al. 2019 |
| pco_3 | - | 98.6 | Y | N | Upham et al. 2019 |
| pco_4 | - | 98.6 | Y | N | Upham et al. 2019 |
| pco_5 | - | 98.6 | Y | N | Upham et al. 2019 |
| pco_6 | - | 98.6 | Y | N | Upham et al. 2019 |
| pco_7 | - | 98.6 | Y | N | Upham et al. 2019 |
| pco_8 | - | 98.6 | Y | N | Upham et al. 2019 |
| pco_9 | - | 98.6 | Y | N | Upham et al. 2019 |
| pco_10 | - | 98.6 | Y | N | Upham et al. 2019 |
| pco_11 | - | 98.6 | Y | N | Upham et al. 2019 |
| pco_12 | - | 98.6 | Y | N | Upham et al. 2019 |
| pco_13 | - | 98.6 | Y | N | Upham et al. 2019 |
| pco_14 | - | 98.6 | Y | N | Upham et al. 2019 |
| pco_15 | - | 98.6 | Y | N | Upham et al. 2019 |
| pco_16 | - | 98.6 | Y | N | Upham et al. 2019 |
| pco_17 | - | 98.6 | Y | N | Upham et al. 2019 |
